# Gene-signatures for early detection of cancers

**DOI:** 10.1101/2023.02.12.528144

**Authors:** Om Prakash, Feroz Khan

## Abstract

**Background:** Gene signatures represents set of molecular modulations in disease genomes or in cells at specific conditions, and are frequently used to classify samples into different groups for better research or clinical treatment. Multiple methods and applications are available in the literature, but powerful ones that can account for early detection of cancer are still lacking.

**Method:** In this article, gene-signatures identified through new in-house algorithm (NCT method) by processing transcriptome data (DEGs extracted from RNA-seq dataset) from population. NCT-Method utilized for processing population dataset, from 28 different human cancer from TCGA & GTEx databases, as empirical background. NCT-score used for optimal clustering of gene-set. The identified gene clusters evaluated through survival analysis. Gene-sets with disease-vs-normal survival plot with logPvalue < 0.05 represented as reliable gene signatures.

**Results:** We applied NCT algorithm to the 28 different cancers, and identified novel gene-signatures as well as inter-relation between different cancers in reference of identified signatures.

**Conclusions:** The algorithm uses population data, and provides validated gene signatures with reliable capacity to discriminate the cancer and normal samples with higher classification performance. The algorithm will be useful to find signature for any RNA-seq data.

## 1. INTRODUCTION

Detection of gene signatures from genomic data is an important topic in medical domain during the last two decades. A “gene signature” can be stated as a single or a group of genes in a cell having a unique pattern of gene expression that is the consequence of either changed biological process or phenotypic behavior. Many computational methods are being used for identification of gene signature out of set of genes. Machine learning techniques are also being involved for multiple myeloma to solving complex problems. Software packages as GeneSpring and other existing R software are known for determination of gene-signature in colorectal cancer through microarray data. Similarly, decision-tree and survival analysis are also known for identification of gene signature for nonsmall-cell lung cancer. So far, a very few attempts has been performed where transition between normal to cancer method has been used for identification of gene signature. In the similar fashion Pareto Evolutionary Algorithm is known for identification of gene-set pattern. Beyond these, some methods have been proposed for optimization of Signatures through multi-objective Genetic Algorithm. The overall performance of these methods are not convincing for early detection of cancer.

The signature gene-set may be a part of differentially expressed gene-set. Statistical analysis is one of the most crucial techniques to determine differentially expressed transcripts across a group of samples versus another group of samples. For RNA-seq data, proper selection of normalization and statistical test are very important, otherwise it might generate wrong p-value for each transcript. Normalized dataset can be obtained from various databases derived from original data at TCGA. Furthermore DEGs are obtained from volcano plot filtering of data. DEGs can be further clustered by using some rule derived from phenotype representation. Determining optimal cluster number of the data is a challenging problem. But it can be overcome by implementing unsupervised clustering followed by identification of significant clusters. Hierarchical clustering along with Silhouette coefficient based evaluation of cluster is useful for such studies. Such strategy support in identification of most appropriate clusters with least error-rate. Of note, in case of the rule derived from phenotype representation problem, the superiority of a solution over other existing solutions & can be produced very easily through the comparison of the scores of their objective functions. Is the rules are derived from population data, then it represents multi-objective optimized solution.

Hence, in this article, we developed an in-house algorithm, Normal to Cancer Transition (NCT) method, of identifying gene signature using population RNA-seq data. Furthermore, population discrimination capacity of gene-signature was also evaluated and justified statistically.

## 2. MATERIALS AND METHOD Population data

RNA expression data, based on tumor and normal samples from the TCGA and the GTEx databases, were collected through Gene Expression Profiling Interactive Analysis web server. RNA expressions for each gene was collected for normal and cancer stage of same tissue. For each gene, median value of RNA expression from population was used. Different population size was available for different cancer type. Initial collection was made for 28 different cancer types.

### Algorithm description

In this article, an algorithm developed for identifying the significant gene-set out of DEGs extracted from RNA-seq expression from a cancer type. Algorithm is based on the feature of transition of normal cell into cancer cell, so named **NC-transition (NCT) algorithm**. The algorithm resultant as gene-set bear significance for discrimination of population between diseased & normal categories. Algorithm was cross-validated with TCGA & GTEx database derived cancer gene expression data. NC-transition algorithm is based on assumptions made for transition (T) of genotype profile from normal to cancer state; which is the very first step in induction on transition from normal to cancer. This transition phase is the indicator for early detection of disease. This transition was picturized in terms of Tscore. Tscore is the function of probabilities of a system of gene-set performing cancer or normal cell status. Description of NCT algorithm has been saved for peer-reviewed evaluation.

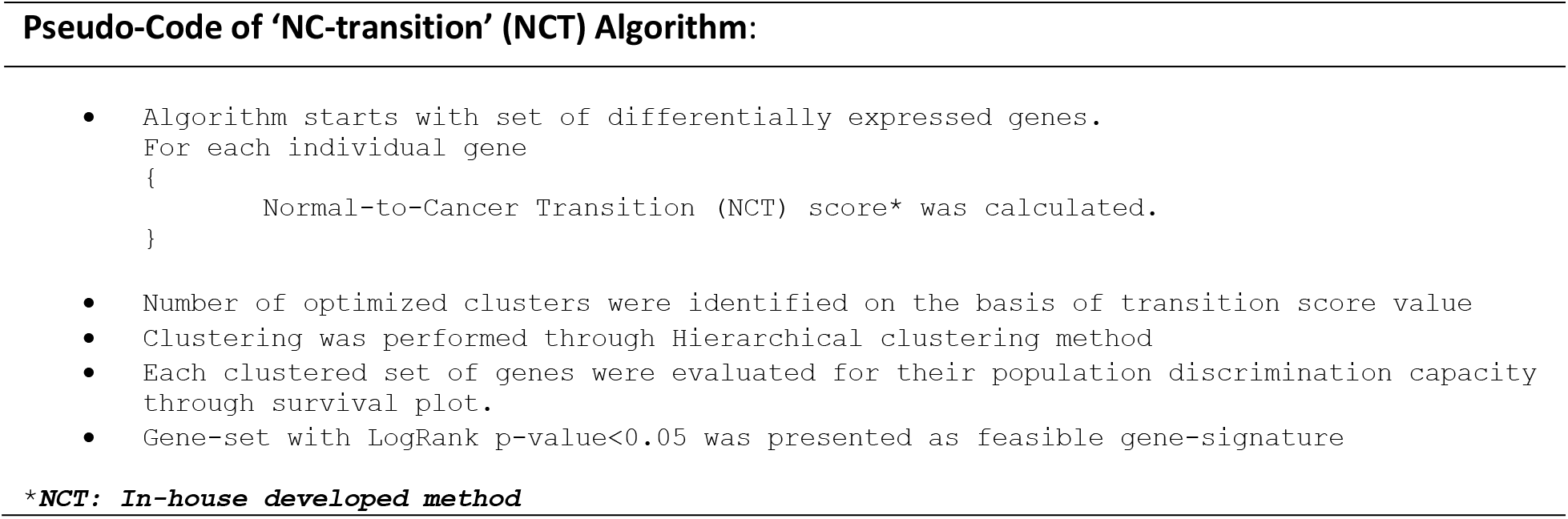

### Finding DEGs

Raw data collected as median value of gene expression from different cancer types. Raw data pre-processed for removal of “all zeros”, “NA values”, and “infinite values”. Then after, data was normalized. Differentially expressed genes (DEGs) identified by volcano plot method. Volcano plot used bi-filtering approach with log-fold change and expression significance value. As a result, a set of differentially expressed genes were identified. Up-regulated genes were ranged with p-value < 0.05 and fold change > 2.0, whereas a down-regulated gene be a gene found between p-value < 0.05 and fold change < 0.5.

### Clustering of DEGs

Unsupervised Hierarchical clustering along with Silhouette method utilized for estimation of optimal cluster number from DEGs. Gene cluster with significant population discrimination was best cluster. The significant cluster was treated as a signature for the respective disease. In present study, we used 28 different cancers for extraction as well as evaluation of gene-signatures.

### Population discrimination evaluation of gene-set

To validate the gene signature, survival plot was estimated for each gene-set cluster. Experimentally validated significant gene-signature identified as survival plot with log-pvalue < 0.05. The whole process was repeated for each gene-set(s) obtained from 28 cancer types. In addition to the gene-set population discrimination evaluation, survival map analysis was also performed for understanding the involvement of same gene-sets in different cancers.

## 3. RESULTS AND DISCUSSUION

Here, we firstly described the source of the population data for 28 different cancers and then performed the experiment and showed the results. We used RNA expression derived from TCGA and the GTEx databases; including tumor and normal samples. Data from each cancer was processed through NCT-algorithm for searching potential gene-signature bearing population discrimination capacity. Total 105 gene-signatures (from 28 cancers) were identified (see supplementary data). Out of 105, only 10 were found to be validated through population data with significant classification capacity. Following are the list of validated gene-signatures from different cancers using population data. Details of all the clustered gene signatures can be accessed from Supplementary table. Description about each validated gene-signature has been given below (Table 1).

**Table 1.**
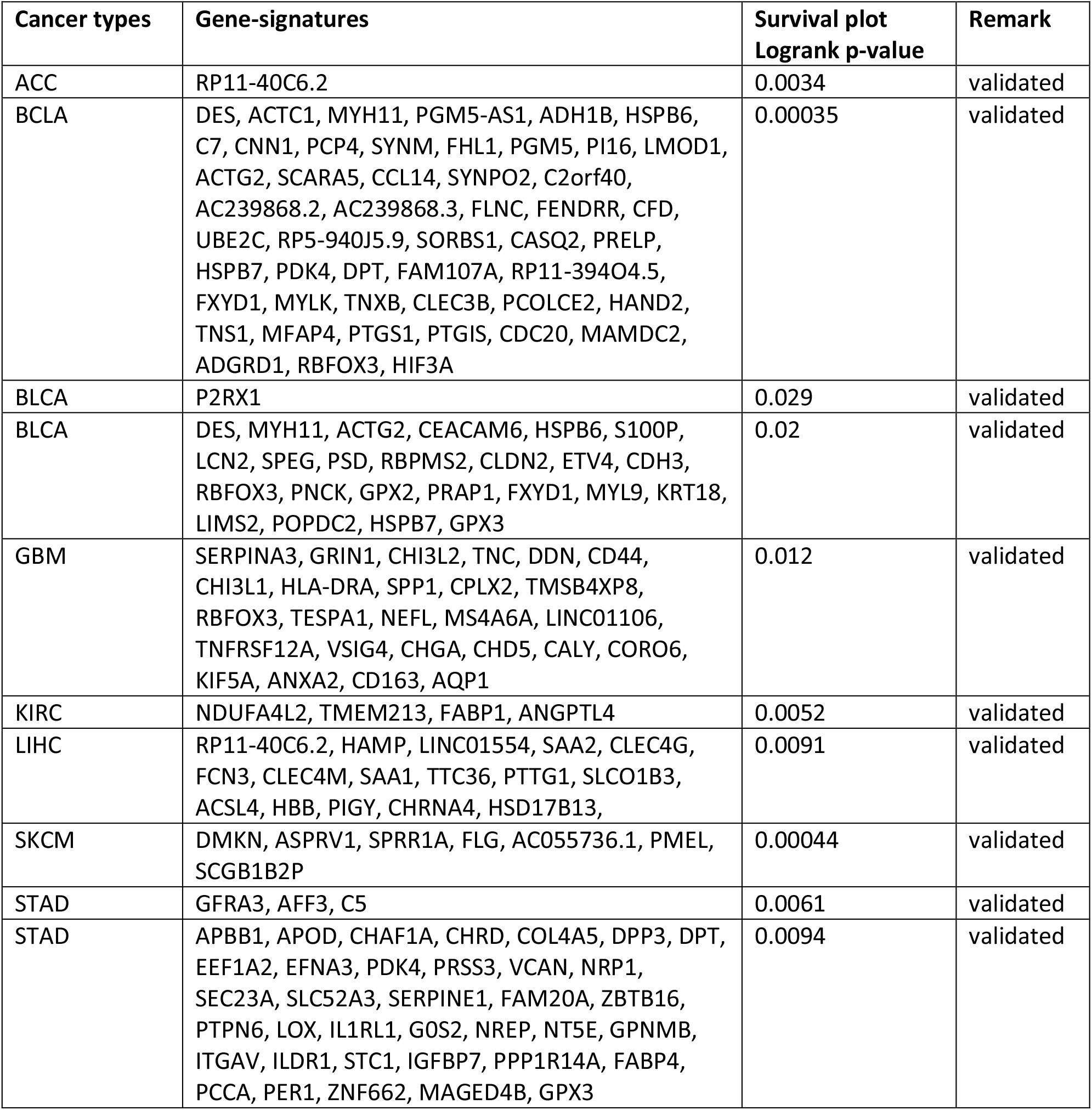
Validated gene-signatures for early detection of cancer.

### 3.1. Gene-signature(s) for Adrenocortical carcinoma (ACC), and their mapping with other cancer types

ACCs are formed in the adrenal cortex. An ACC may be of two types: (i) making more hormone than normal, (ii) has no effect on hormone production. First category often makes too much of the hormones cortisol, aldosterone, testosterone, or estrogen. In present study included 128 Normal and 77 Tumor samples of ACC. 08 optimal clusters formed from the DEGs; out of which only one cluster (Cluster-1) was found to be efficient for population discrimination with Logrank p-value of 0.0034. The validated cluster gene-set contained only one gene, named as ‘RP11-40C6.2’; identified through optimal clustering of ACC-DEGs, performed significantly at survival plot and box-plot. RP11-40C6.2 is a ribosomal protein S29 (RPS29) pseudogene, encoding a ribosomal protein. It belongs to the S14P family of ribosomal proteins; which contains a C2-C2 zinc finger-like domain for enhancing the tumor suppressor activity of Ras-related protein 1A (KREV1). Variable expression of this gene was observed in colorectal cancers, but by comparing the normal tissues, no correlation was identified between the extent of expression and intensity disease. KREV1 (UniProtKB ID: P62834) is known to induces morphological reversion of a cell line which are transformed by a Ras oncogene. It also counteracts the mitogenic function of Ras, because of its ability to interact with both Ras GAPs and RAF. It also regulates KRIT1 localization to microtubules and membranes, with support of ITGB1BP1. It also participates in neurite outgrowth by induction of nerve growth factor (NGF). It also participates in the regulation of formation of blood vessels as well as establishment of basal endothelial barrier function. It may also be involved in the regulation of the vascular endothelial growth factor receptor KDR expression at endothelial cell-cell junctions. These interpretations suggests that during the transition of normal to cancer cell (possibly at the early stage of cancer development), the cell-to-cell junction get affected [1, 2]. Survival map was drawn on the basis of performance of ‘RP11-40C6.2’ through log10 of Hazard ratio. It was observed that the identified gene-signature may show similar discrimination in population of KICH & PCPG cancer types (Figure 1).

**Figure 1.**
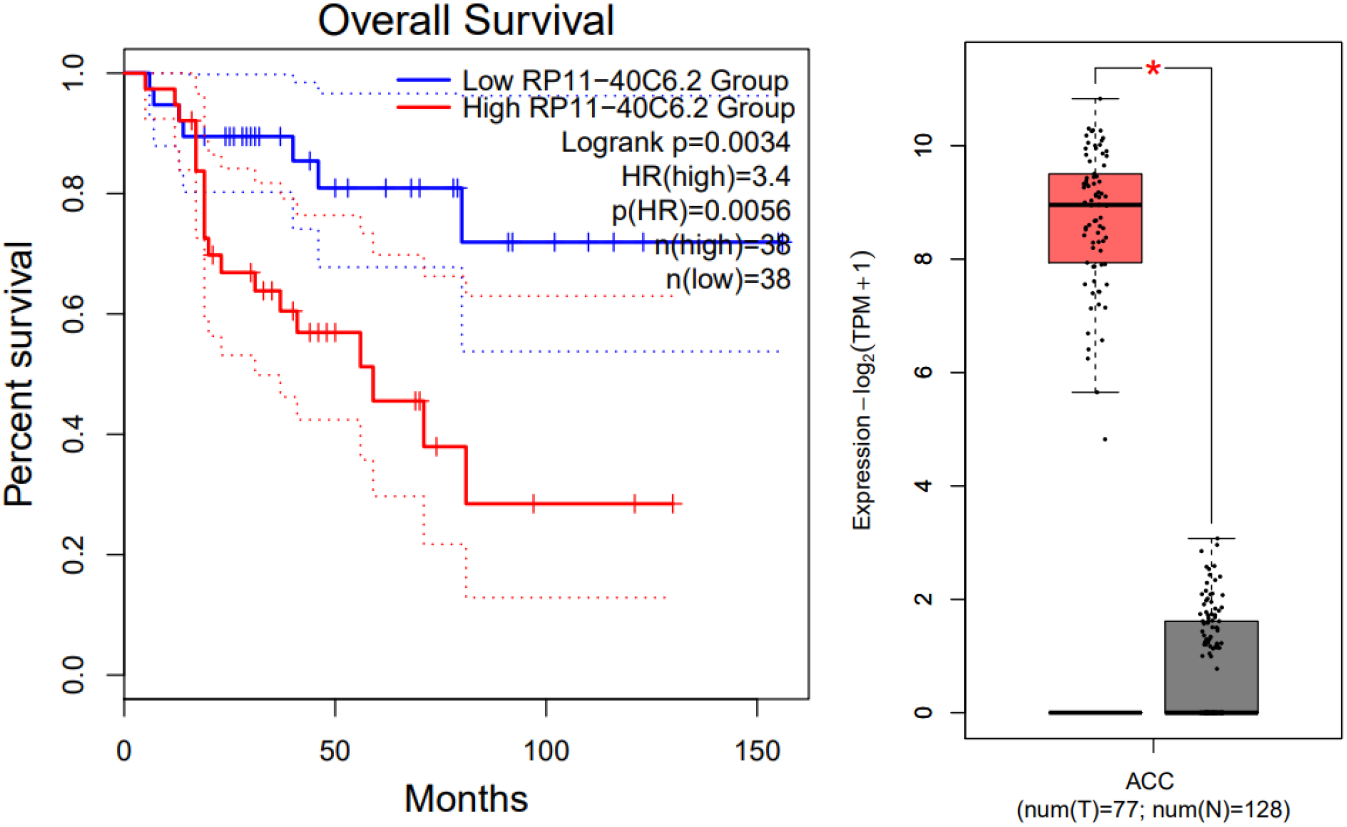
Cluster-1 gene-set (‘RP11-40C6.2’) as significant gene signature, identified through optimal clustering of ACC-DEGs, performed significantly at survival plot and box-plot.

### 3.2. Gene-signature(s) for Bladder Urothelial Carcinoma (BCLA), and their mapping with other cancer types

BCLA is the cancer of urinary bladder. In present study, 28 Normal and 404 Tumor samples of BCLA were considered. Optimal clustering resulted into two groups. Two clusters were found to be efficient for population discrimination with Logrank p-value of 0.00035 (cluster-1) and 0.029 (cluster-2). **Cluster-1**, gene-set co-expression suggested for molecular function of Actin binding (GO: 0003779) and Cytoskeleton protein binding (GO: 0008092), which are basically related with pathways of smooth muscle contraction (HSA-445355 and HSA-397014). Prior study by Hwang et al, 2020, mentioned involvement of smooth muscle in Bladder Cancer [3]. **Cluster-2**, found to be linked with apoptosis, where increasing the intracellular concentration of calcium in the presence of ATP, leading to programmed cell death. Prior study by Dillard et al 2021, demonstrated for similar type of observation [4]. Survival map, drawn on the basis of performance of cluster-1&2 through log10 of Hazard ratio, suggested similar capacity of population discrimination in KIRP & SKCM cancer types (Figure 2).

**Figure 2.**
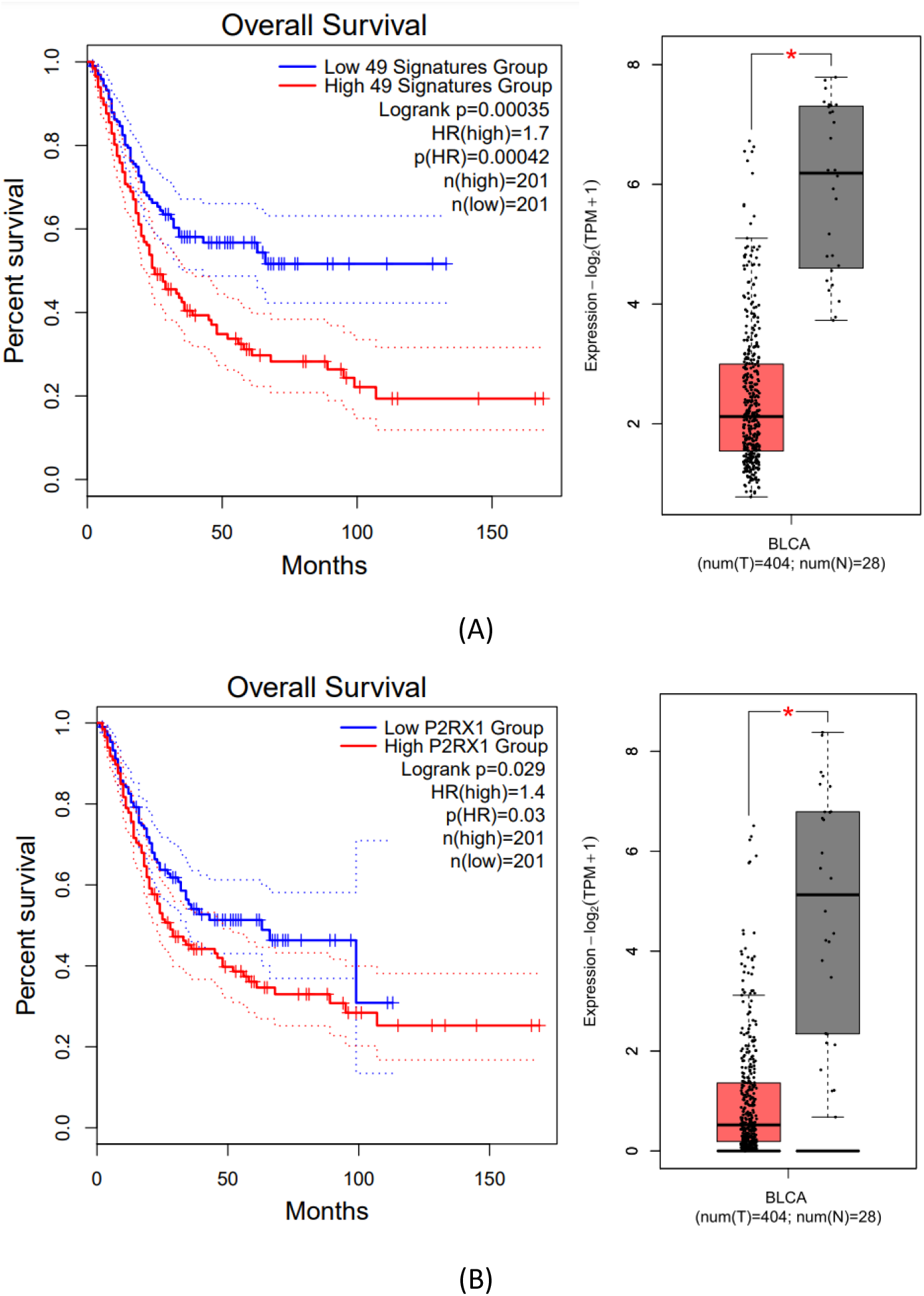
Cluster-1(A) &2(B) gene-set as significant gene signature, identified through optimal clustering of BCLA-DEGs, showed significant survival plot and box-plots.

### 3.3. Gene-signature(s) for Colon adenocarcinoma (COAD), and their mapping with other cancer types

COAD is the cancer of Colon. In present study, 349 Normal and 275 Tumor samples of COAD were considered. Optimal clustering approved for 04 optimal clusters out from the DEGs. But the most efficient for population discrimination was with Logrank p-value of 0.02 (cluster-2). Cluster-2, gene-set co-expression suggested for molecular function at Cell-cell junction (GO:0005911); basically, related with pathways of smooth muscle contraction (HSA-445355 and HSA-397014). Prior study by Lee & Daar 2009, also performed in similar fashion, describing EphrinB reverse signaling in cell-cell adhesion [5] (Figure 3).

**Figure 3.**
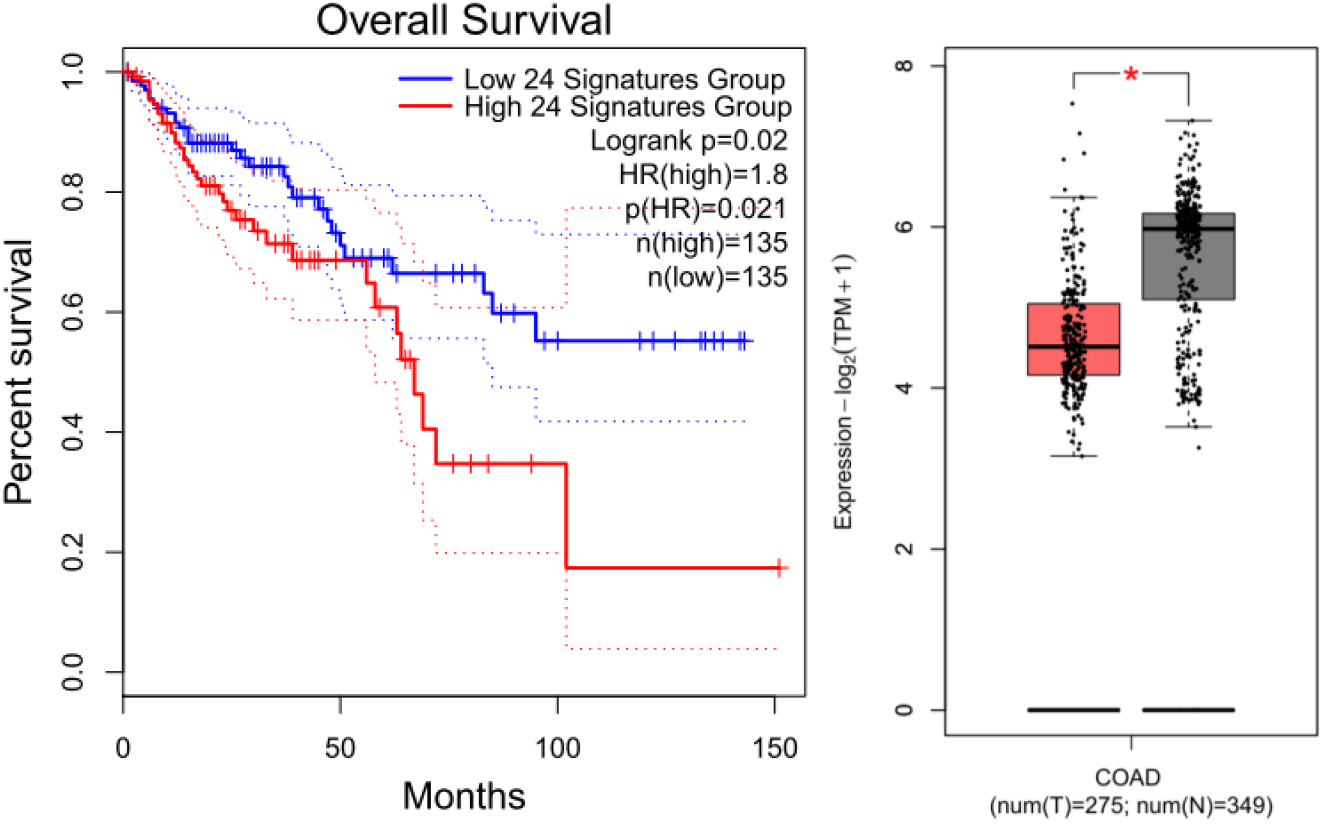
Cluster-2 gene-set as significant gene signature, identified through optimal clustering of COAD-DEGs, showed significant survival plot and box-plots.

### 3.4. Gene-signature(s) for (GBM), and their mapping with other cancer types

Glioblastoma (GBM) is **an aggressive type of cancer that can occur in the brain or spinal cord**. In present study, 207 Normal and 163 Tumor samples of GBM were considered. Optimal clustering approved for 03 clusters out from the DEGs. One cluster was found to be efficient for population discrimination with Logrank p-value of 0.012 (cluster-1). Cluster-1, gene-set co-expression indicated towards the impact localization in extra-cellular region, plasma membrane, cytoskeleton and connective tissue. Mohiuddin & Wakimoto, 2021 also suggested for involvement of Extracellular matrix in glioblastoma for possible opportunities of therapeutic approaches [6] (Figure 4).

**Figure 4.**
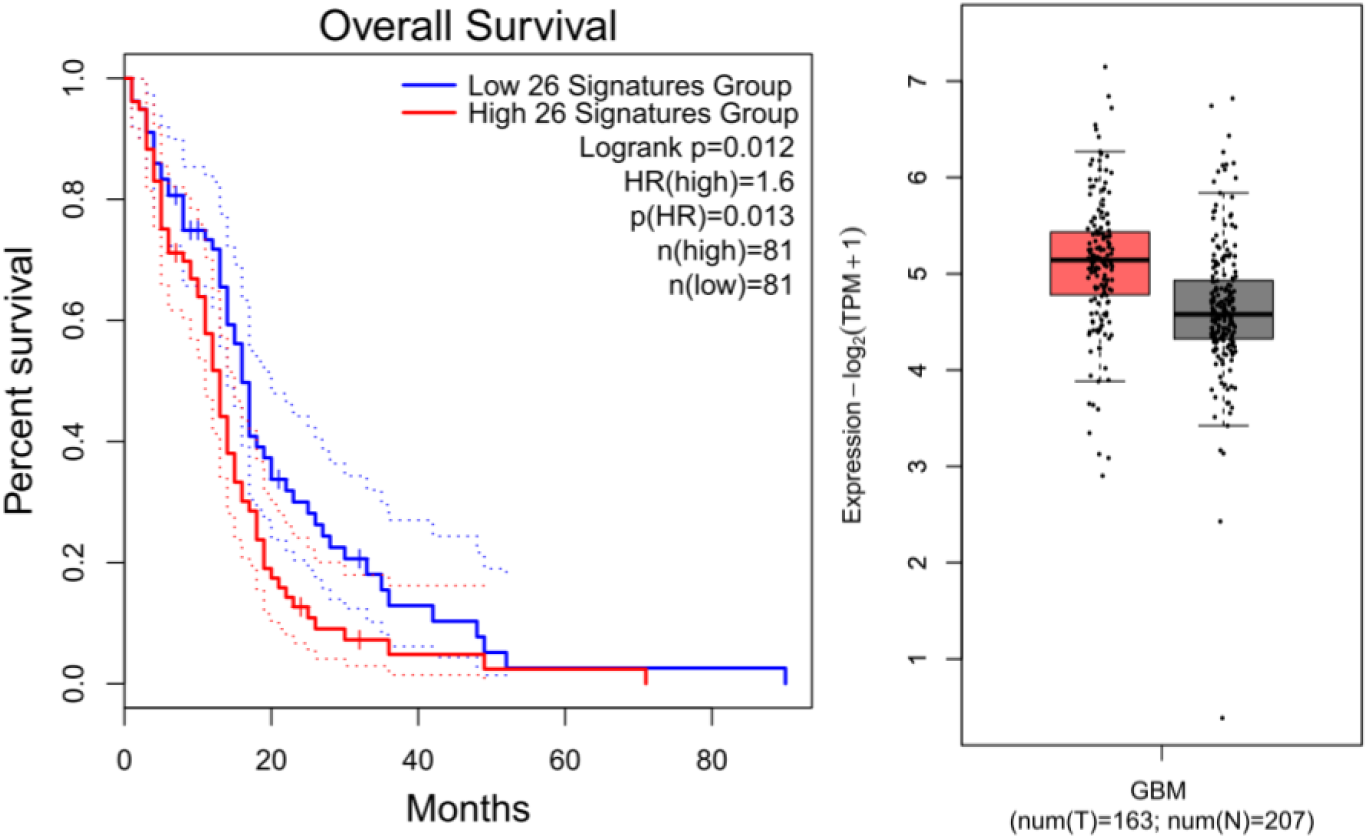
Cluster-1 gene-set as significant gene signature, identified through optimal clustering of GBM-DEGs, showed significant survival plot and box-plots.

### 3.5. Gene-signature(s) for Kidney (KIRC), and their mapping with other cancer types

KIRC is the cancer of Kidney. In present study, 100 Normal and 523 Tumor samples of KIRC were considered. Optimal clustering resulted into two clusters. But only one was found to be efficient for population discrimination with Logrank p-value of 0.0052 (cluster-2). Cluster-2, gene-set expression was found to be linked with PPAR signaling pathway [6] (Figure 5).

**Figure 5.**
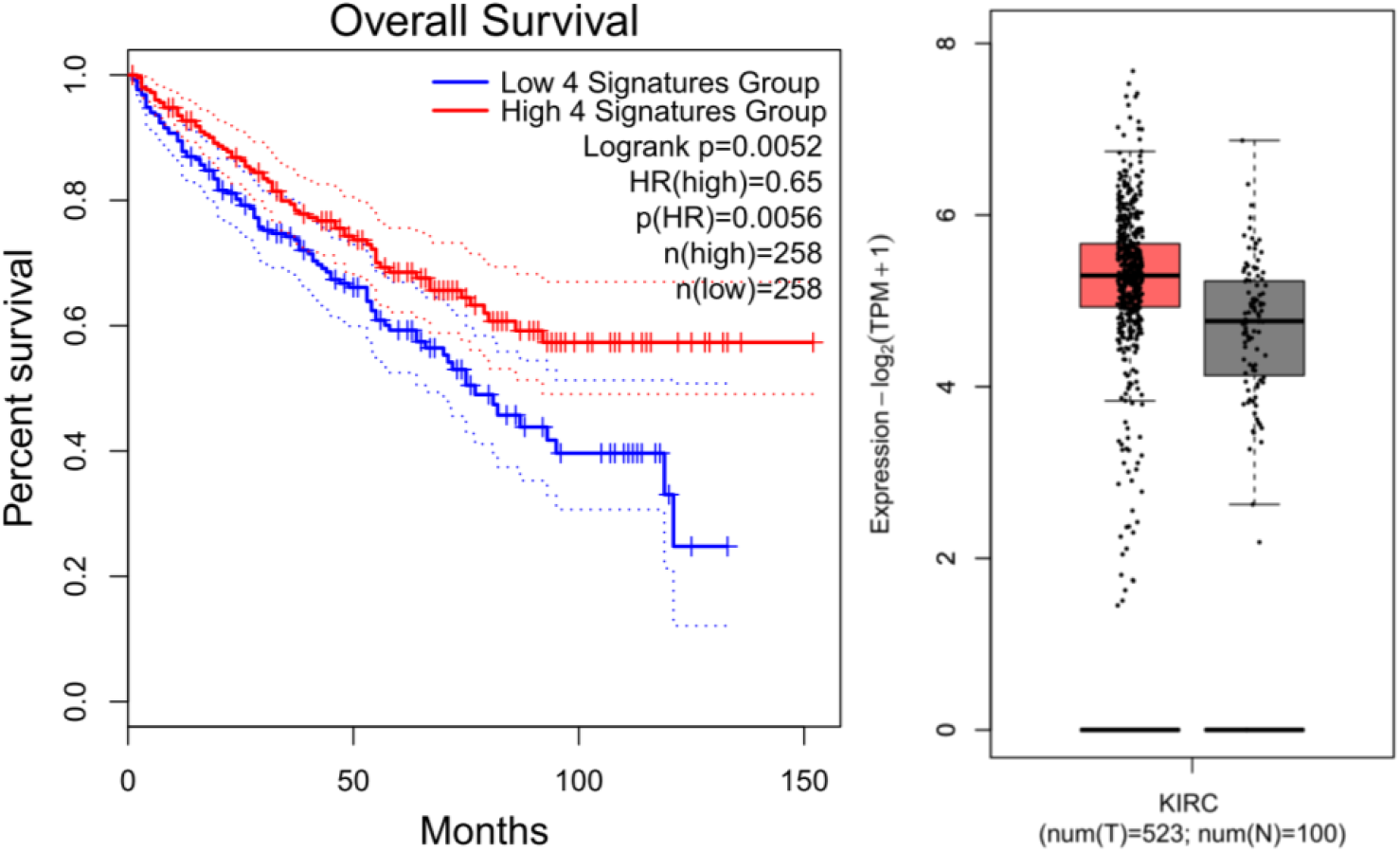
Cluster-2 gene-set as significant gene signature, identified through optimal clustering of KIRC-DEGs, showed significant survival plot and box-plots.

### 3.6. Gene-signature(s) for Liver hepatocellular carcinoma (LIHC), and their mapping with other cancer types

LIHC is the cancer of Liver. In present study, 160 Normal and 369 Tumor samples of LIHC were considered. Optimal clustering resulted into03 clusters, but only one was efficient for population discrimination with Logrank p-value of 0.0091 (cluster-1). Cluster-1, the gene-set was found to be expressed in Liver, and found to be related with HDL, Amyloid protein and hemolytic anemia. It was also found to be linked with metabolism of Folate & Vita-12 [8, 9, 10] (Figure 6).

**Figure 6.**
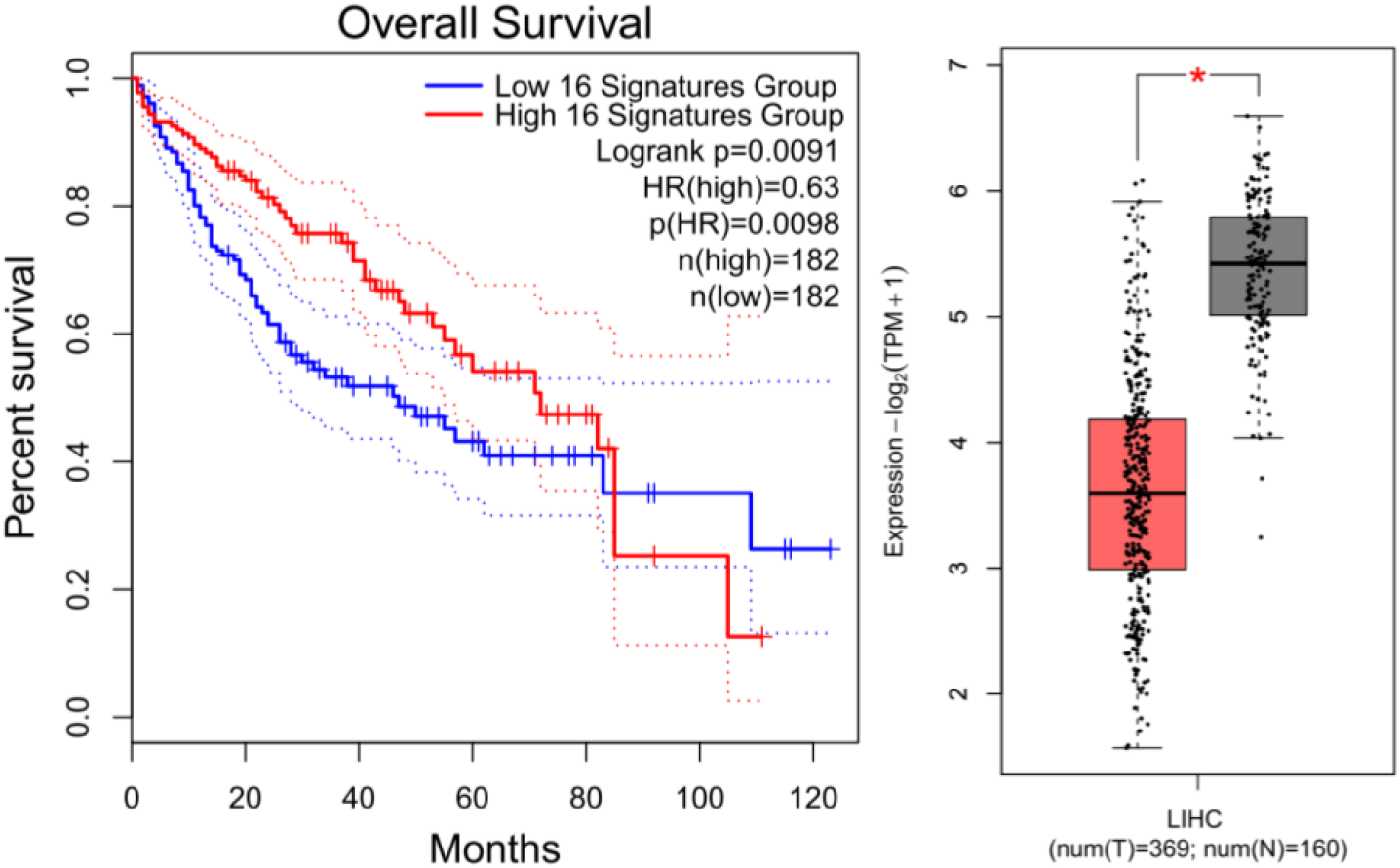
Cluster-1 gene-set as significant gene signature, identified through optimal clustering of LIHC-DEGs, showed significant survival plot and box-plots.

### 3.7. Gene-signature(s) for Skin Cutaneous Melanoma (SKCM), and their mapping with other cancer types

SKCM is the cancer of Skin. In present study, 558 Normal and 461 Tumor samples of SKCM were considered. Optimal clustering resulted into two clusters, but only one to be efficient for population discrimination with Logrank p-value of 0.00044 (cluster-2). Cluster-2: The gene-set co-expression was found to be directly linked with structural constituents of skin epidermis (GO: 0030280), and also tell about the integrity of the epidermis [11] (Figure 7).

**Figure 7.**
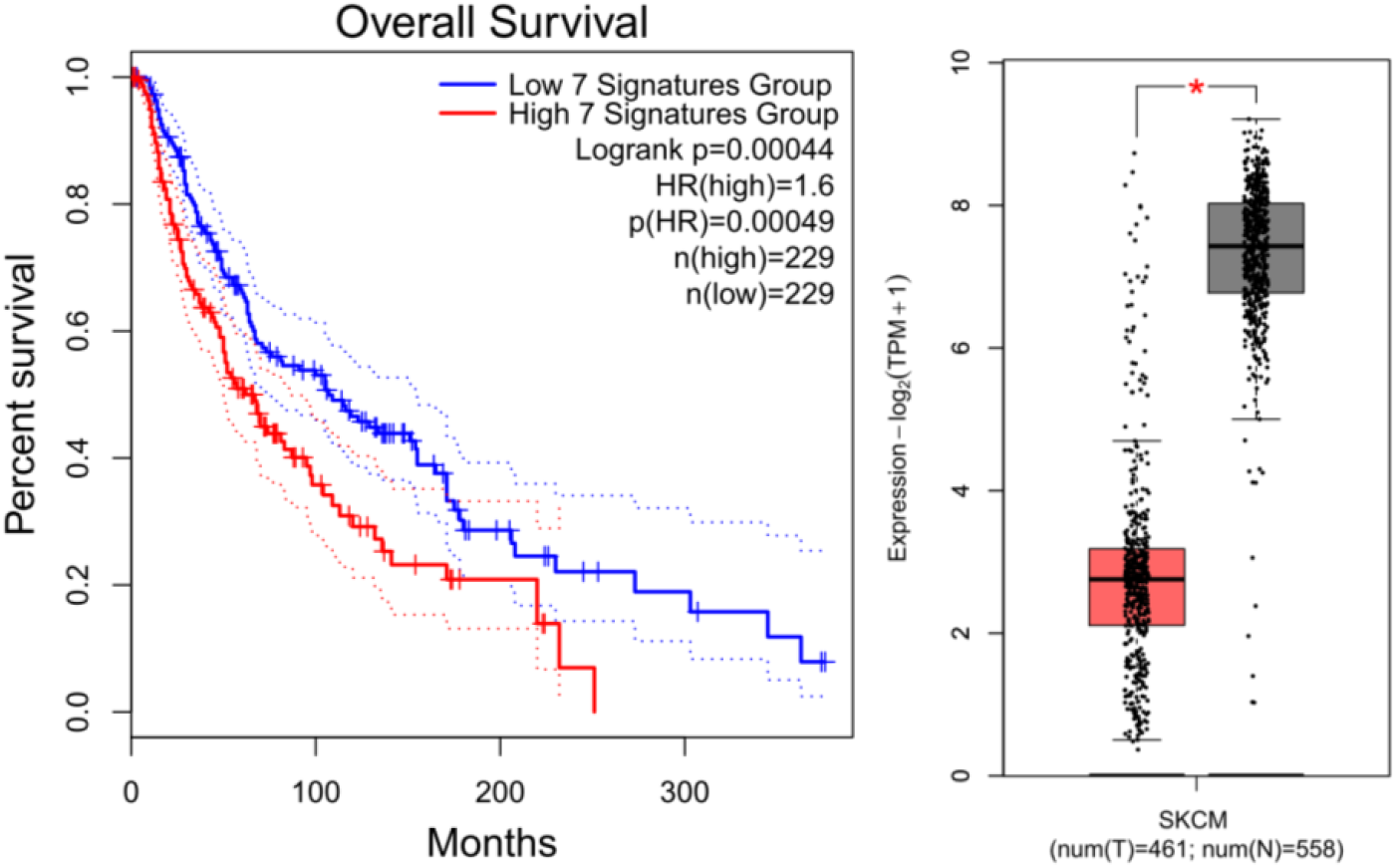
Cluster-2 gene-set as significant gene signature, identified through optimal clustering of SKCM-DEGs, showed significant survival plot and box-plots.

### 3.8. Gene-signature(s) for Stomach Adenocarcinoma (STAD), and their mapping with other cancer types

STAD is the cancer of stomach. In present study, 211 Normal and 408 Tumor samples of STAD were considered. Optimal clustering resulted into two clusters. Both clusters were found to be efficient for population discrimination with Logrank p-value of 0.0061 (cluster-1) and 0.0094 (cluster-2). Cluster-1, gene-set co-expression did not suggested for any cumulative function. While, Cluster-2: gene-set co-expression suggest for their existence for extracellular performance, collagen formation, matrix metalloproteinases, MFAP2 (PMID: 32054827), IL-1, and CD-73 [12, 13, 14, 15] (Figure 8, 9).

**Figure 8.**
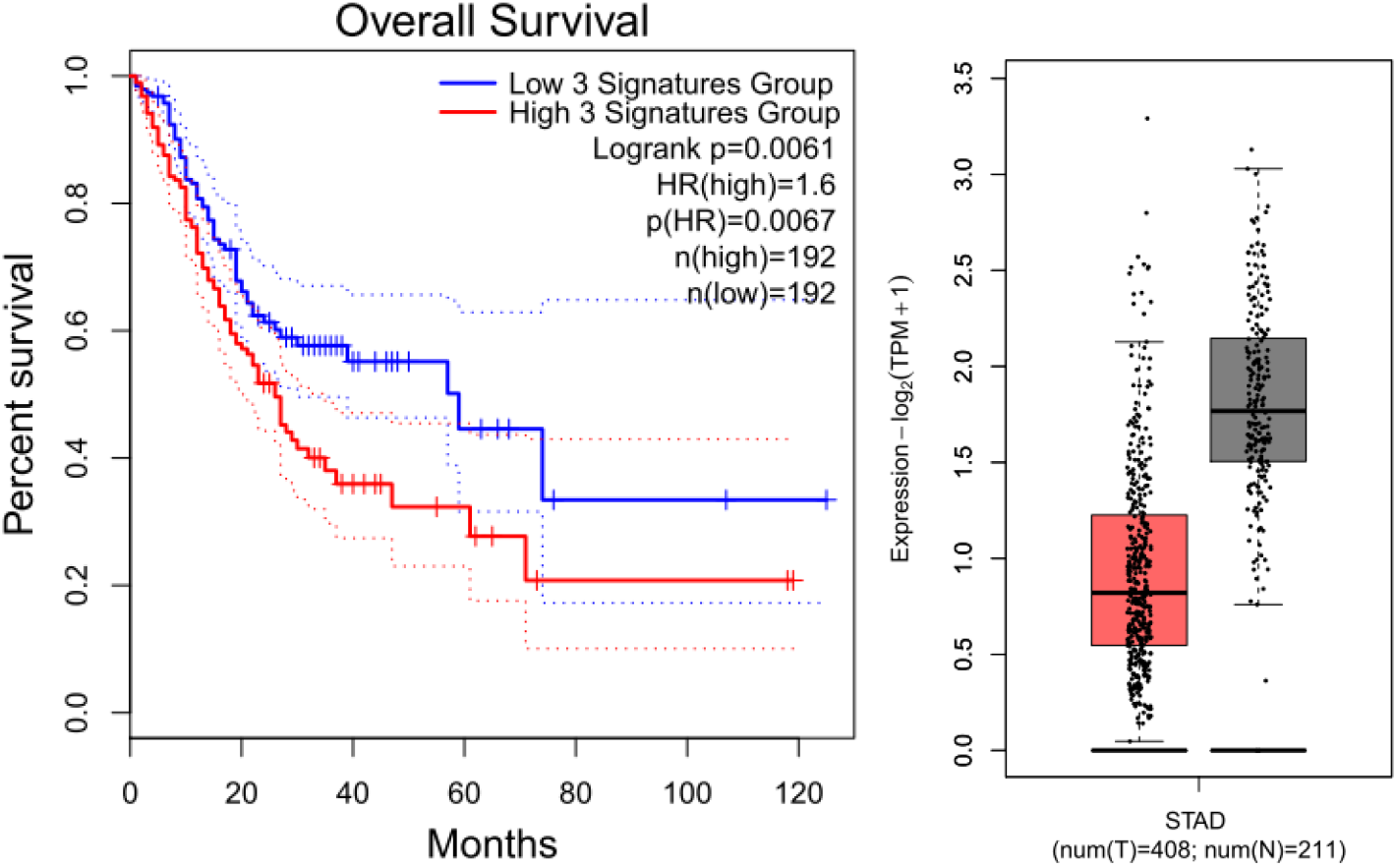
Cluster-1 gene-set as significant gene signature, identified through optimal clustering of STAD-DEGs, showed significant survival plot and box-plots.

**Figure 9.**
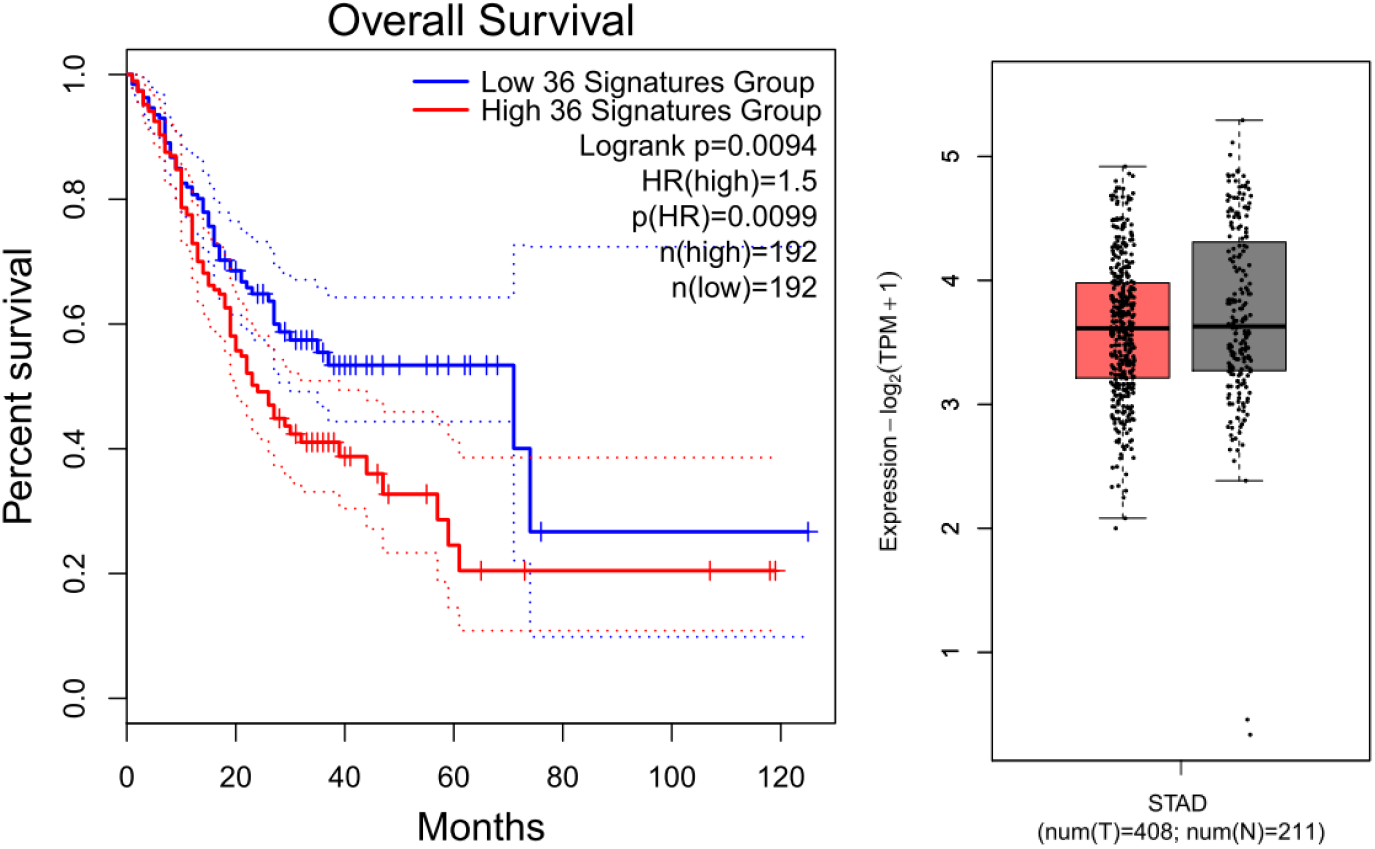
Cluster-2 gene-set as significant gene signature, identified through optimal clustering of STAD-DEGs, showed significant survival plot and box-plots.

## CONCLUSION

Although there are many bioinformatics approaches available for the gene signature identification, the gene signature identification for early detection need improvement; Therefore, in this article, we developed a new framework of identifying gene signature using NC-transition (NCT-algorithm) through RNA-seq data, followed by experimental validation through survival plotting. In this regard, we conducted this study with DEGs from population data for a specific cancer. Next, we applied ANOVA method to find the differentially expressed genes consisting of up-regulated and down-regulated genes. Thereafter, we applied Silhouette method on these differentially expressed genes to estimate the optimal cluster size, and then applied hierarchical clustering through the optimal cluster size. The best cluster was obtained through computing the survival plot for each cluster. The best clusters were treated as a signature for the respective disease. In this work, we used 28 different cancers from TGCA database. The study concluded into 10 identified gene signatures. The identified signatures might be helpful for early diagnosis of the cancer. Finally, our method is useful to identify gene signature for any RNA-seq or similar kind of data.

## Acknowledgement

Authors are thankful to the Director, CSIR-Central Institute of Medicinal & Aromatic Plants (CIMAP), Lucknow, India for infrastructure & research facilities support. Author OP is thankful to the Indian Council of Medical Research (ICMR), New Delhi, India for financial support through RA fellowship (Award letter no. BMI/11(12)/2020, dated: 04/02/2021). The CSIR-CIMAP publication number of this manuscript is CIMAP/PUB/ 2023/20.

## Conflict of interest

There is no conflict of interest.

## Supplementary data

**Table S1.** List of cancers considered in the study

| TCGA | Detail |
| --- | --- |
| ACC | Adrenocortical carcinoma |
| BLCA | Bladder Urothelial Carcinoma |
| BRCA | Breast invasive carcinoma |
| CESC | Cervical squamous cell carcinoma and endocervical adenocarcinoma |
| CHOL | Cholangio carcinoma |
| COAD | Colon adenocarcinoma |
| DLBC | Lymphoid Neoplasm Diffuse Large B-cell Lymphoma |
| ESCA | Esophageal carcinoma |
| GBM | Glioblastoma multiforme |
| HNSC | Head and Neck squamous cell carcinoma |
| KICH | Kidney Chromophobe |
| KIRC | Kidney renal clear cell carcinoma |
| KIRP | Kidney renal papillary cell carcinoma |
| LAML | Acute Myeloid Leukemia |
| LGG | Brain Lower Grade Glioma |
| LIHC | Liver hepatocellular carcinoma |
| LUAD | Lung adenocarcinoma |
| LUSC | Lung squamous cell carcinoma |
| MESO | Mesothelioma |
| OV | Ovarian serous cystadenocarcinoma |
| PAAD | Pancreatic adenocarcinoma |
| PCPG | Pheochromocytoma and Paraganglioma |
| PRAD | Prostate adenocarcinoma |
| READ | Rectum adenocarcinoma |
| SARC | Sarcoma |
| SKCM | Skin Cutaneous Melanoma |
| STAD | Stomach adenocarcinoma |
| TGCT | Testicular Germ Cell Tumors |
| THCA | Thyroid carcinoma |
| THYM | Thymoma |
| UCEC | Uterine Corpus Endometrial Carcinoma |
| UCS | Uterine Carcinosarcoma |
| UVM | Uveal Melanoma |

**Table S2.**
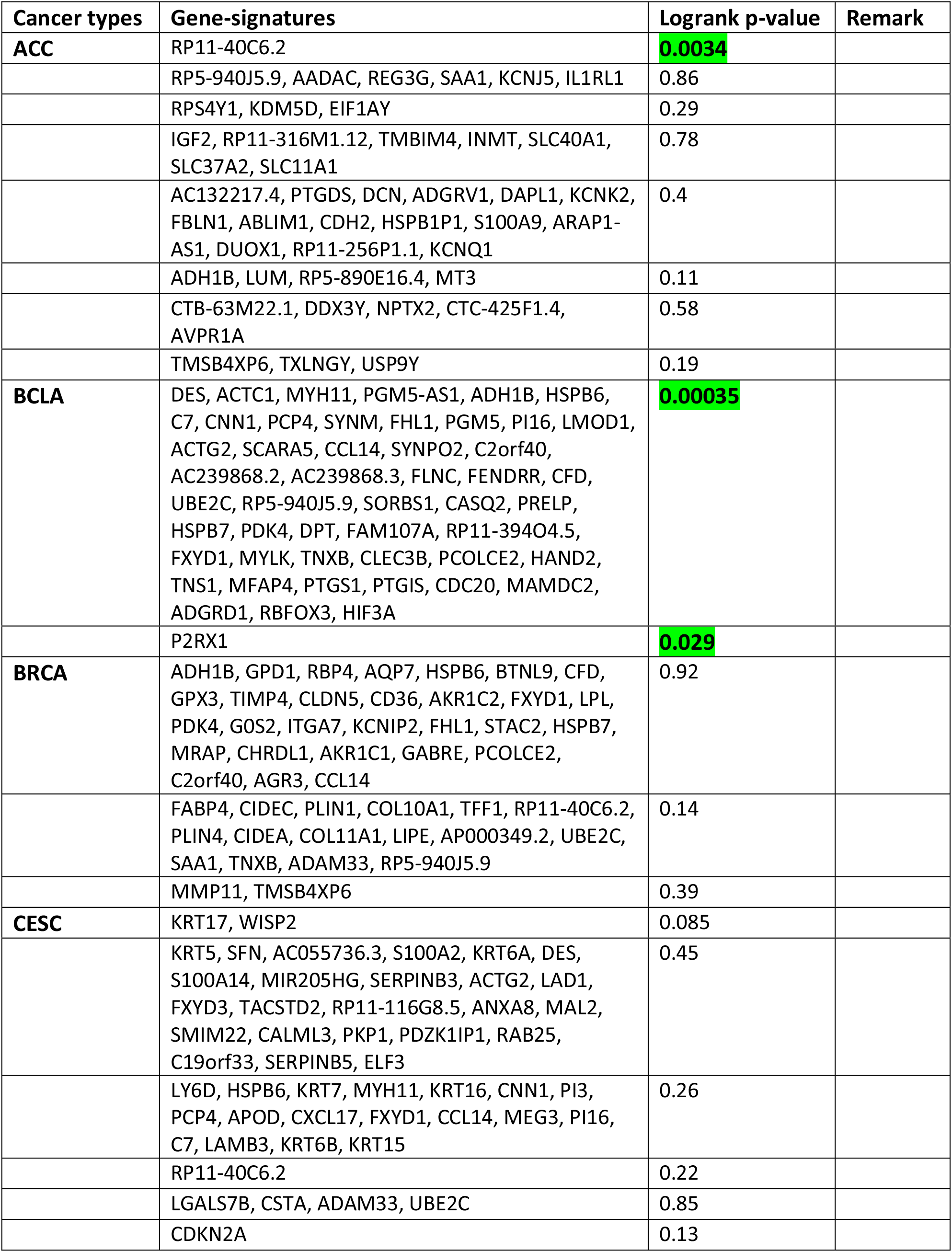
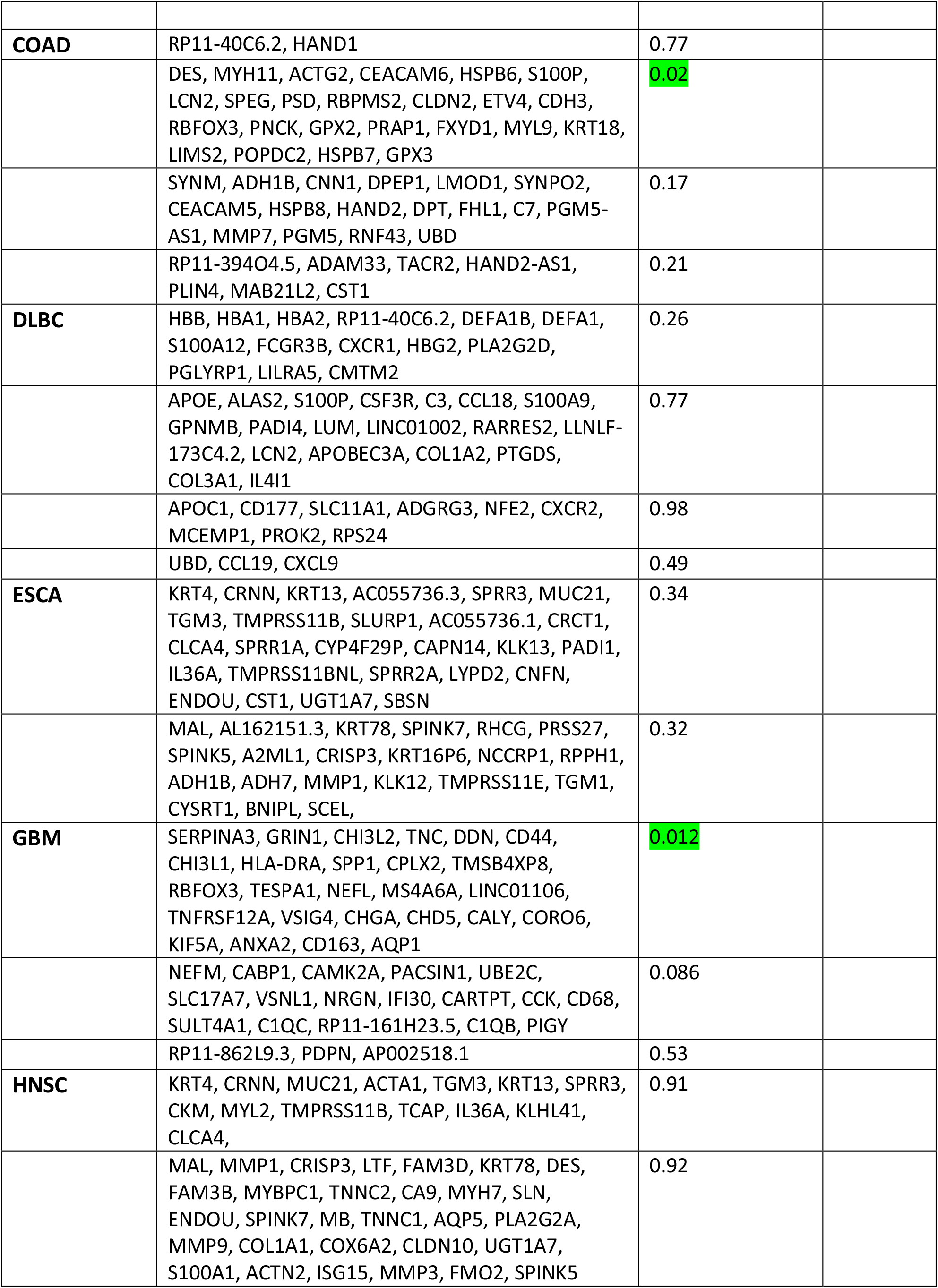
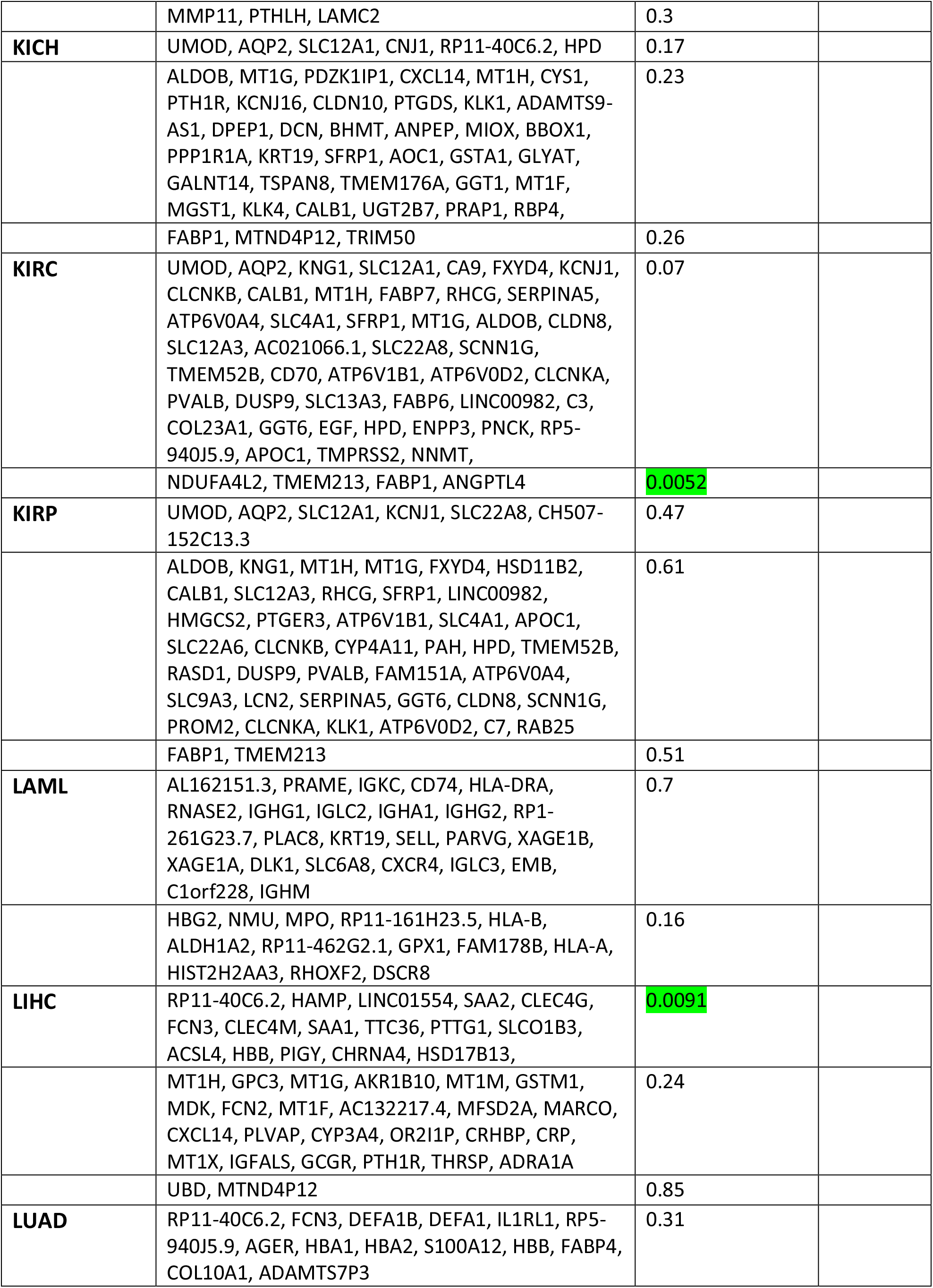
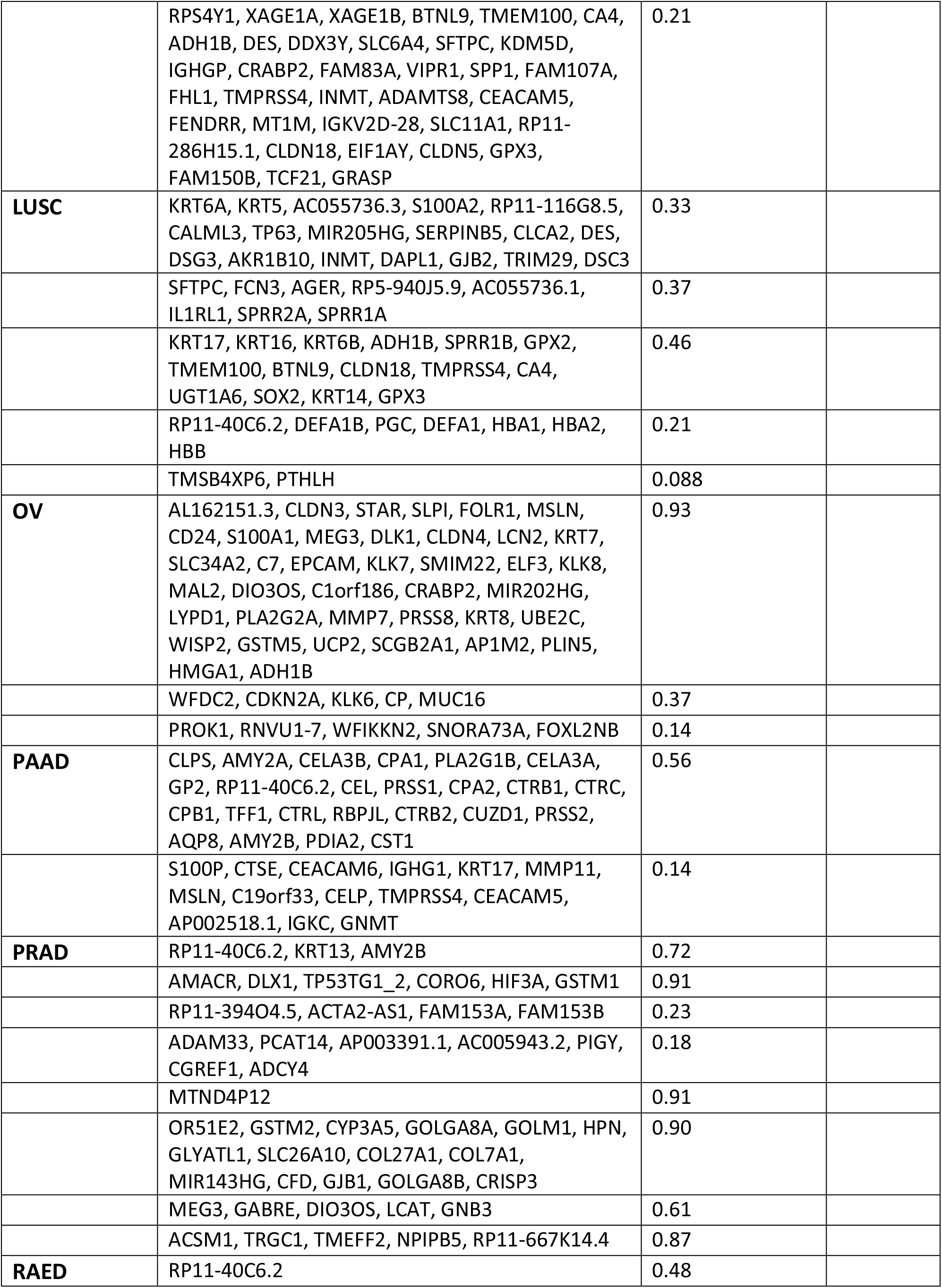
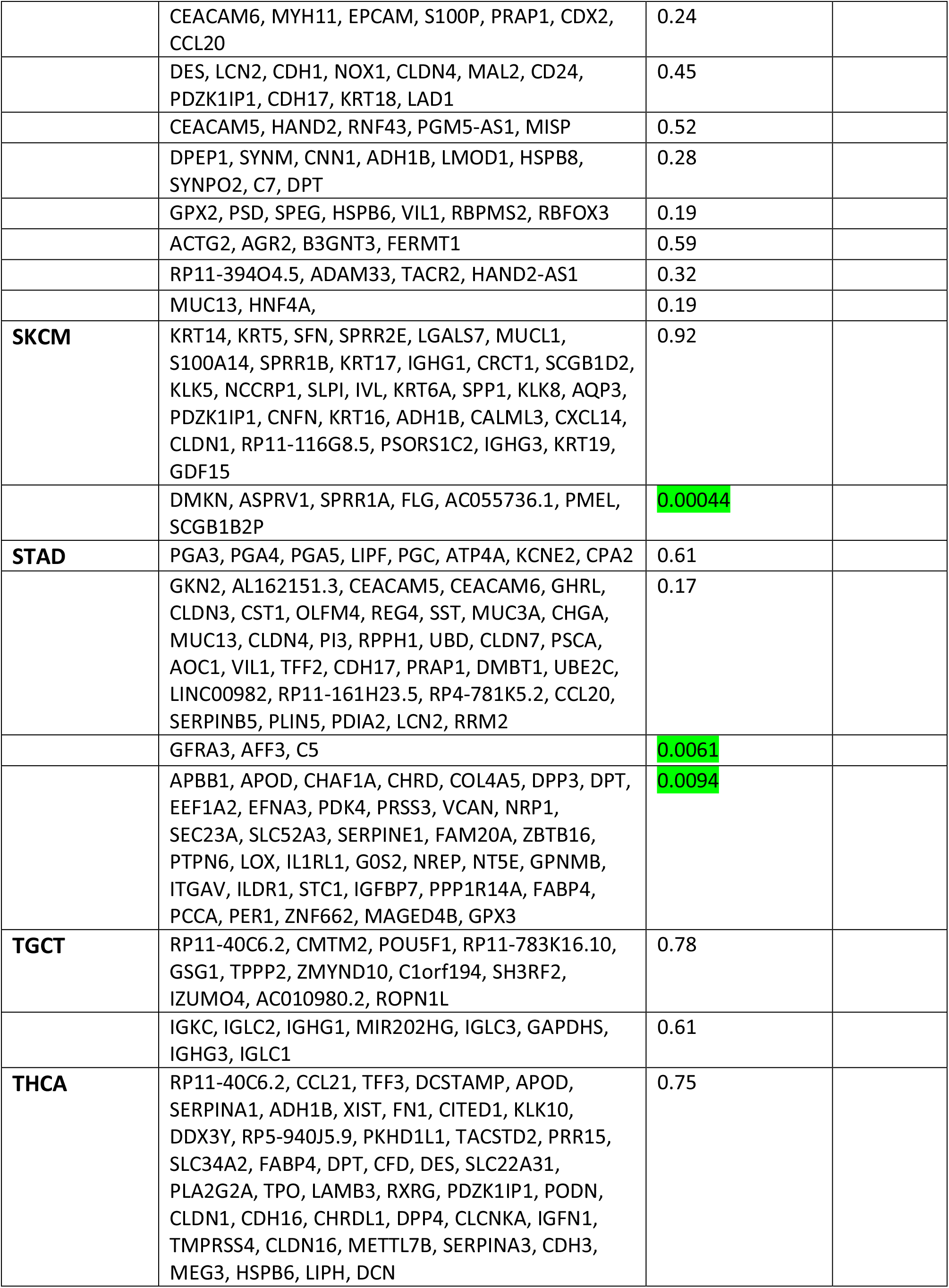
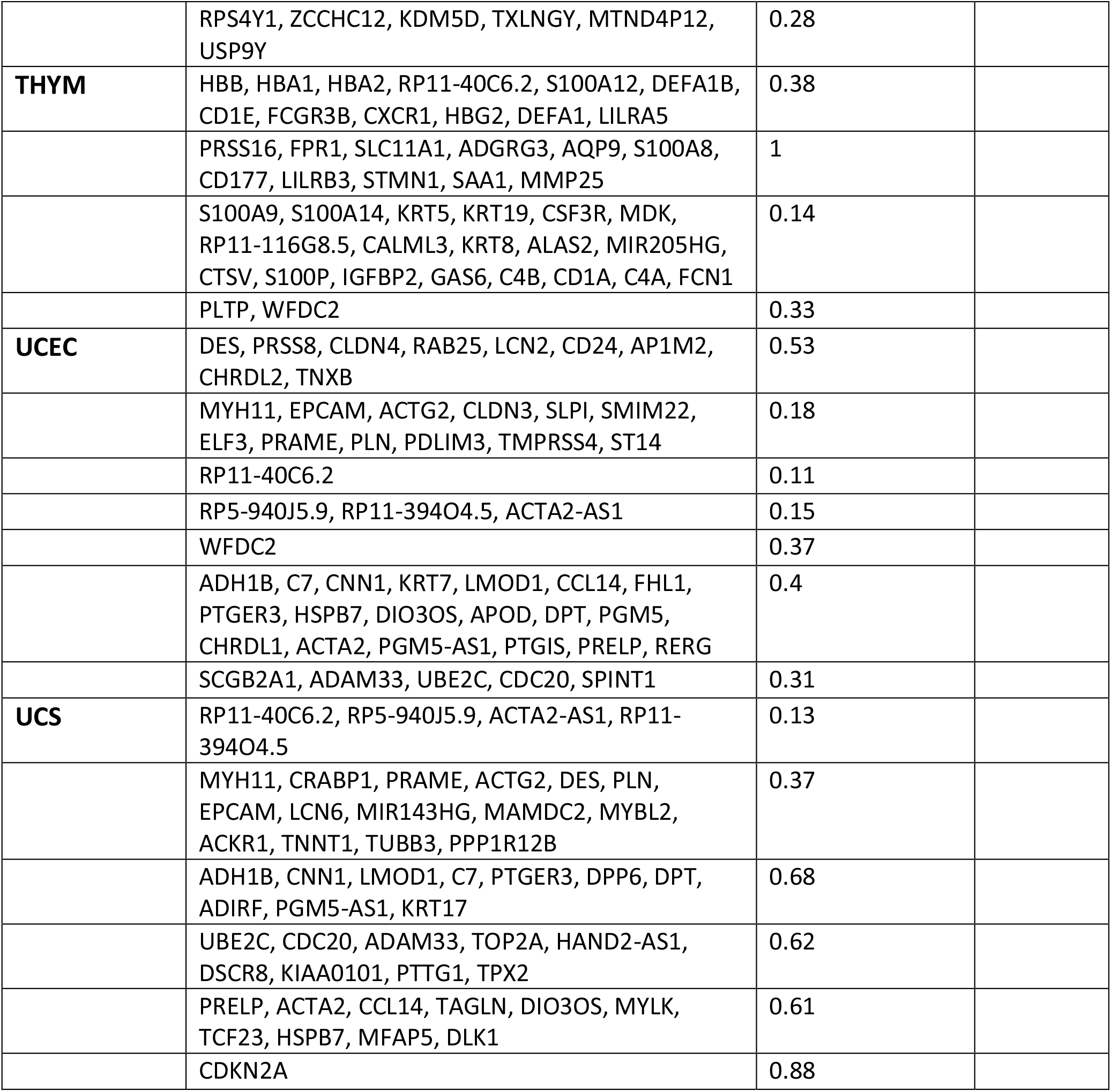
Gene-signatures Identified from 20 cancers

## Notes

### Competing Interest Statement

The authors have declared no competing interest.

